# Building new compartments for unconventional protein secretion from the early and late Golgi membranes

**DOI:** 10.1101/2021.10.22.465504

**Authors:** Amy J. Curwin, Nathalie Brouwers, Akihiko Nakano, Kazuo Kurokawa, Vivek Malhotra

**Affiliations:** Centre for Genomic Regulation, The Barcelona Institute of Science and Technology, 08003 Barcelona, Spain; Universitat Pompeu Fabra (UPF), 08002 Barcelona, Spain; ICREA, 08010 Barcelona, Spain; Live Cell Super-Resolution Imaging Research Team, RIKEN Center for Advanced Photonics, Wako, Saitama, Japan

## Abstract

CUPS, a compartment for unconventional secretion of signal sequence lacking proteins, is built during starvation. CUPS, lacking the Golgi specific glycosyltransferases, form by COPI independent extraction of membranes from the early Golgi cisterna, require PI4P for their biogenesis and PI3P for stability. We now show that a PI4P effector Drs2 of the trans-Golgi network, relocates to a new compartment monikered TCUPS because it touches CUPS. Although localized to TCUPS, Drs2 is required for CUPS formation specifically by interacting with Rcy1, and this process is essential for unconventional secretion. Visualizing cells by 4D SCLIM technology revealed that tubules emanating from TCUPS are often collared by CUPS and severed. Incidentally, while CUPS are stable, TCUPS are vesiculated at late stages of starvation. This mirrors the dynamics of the early and late Golgi during conventional protein secretion. TCUPS and CUPS thus emerge as the functional equivalent of early and late Golgi of the conventional secretory pathway, thus representing key compartments in unconventional secretion.

## Introduction

The problem of how proteins that cannot enter the endoplasmic reticulum (ER)-Golgi pathway of secretion are released to the extracellular space remains a fascinating challenge. This is an important issue because cells secrete proteins like fibroblast growth factor (FGF) 2, Interleukin (IL)-1ß, Acyl CoA binding protein (Acb1/Diazepam binding inhibitor), Superoxide dismutase (SOD) 1, and tissue transglutaminase, which have important physiological roles in the extracellular space, particularly under conditions of stress. The prevailing schemes for the secretion of this class of cytoplasmic proteins include: involvement of an intracellular compartment, such as CUPS (compartment for unconventional protein secretion), for proteins like Acb1, SOD1 and many other antioxidants, or secretory endosomes or lysosomes, in the case of fatty acid binding protein 4 (FABP4), or direct translocation across the plasma membrane for FGF2 (Bruns et al., 2011; Cruz-Garcia et al., 2020; Schäfer et al., 2004; Villeneuve et al., 2017). IL-1ß is reported to use multiple routes of export such as translocation directly across the plasma membrane via a pore created by Gasdermin D, translocation by conventional cargo receptor protein TMED10 into the ER-Golgi intermediate compartment (ERGIC) prior to its release from cells, and by pyroptosis (He et al., 2015; Liu et al., 2016; Zhang et al., 2020).

There might indeed be different routes for this mode of transport, but we have focussed on the pathway for secretion of Acb1 and Sod1. The export of these proteins has the following essential requirements. 1, Their secretion is triggered upon carbon and nitrogen starvation, and growth in potassium acetate; 2, intracellular production of reactive oxygen species (ROS); 3, the need of a peripherally Golgi/ER exit site localized protein called Grh1 (GORASPs in mammals); and 4, a compartment called CUPS (Bruns et al., 2011; Cruz-Garcia et al., 2014, 2020; Curwin et al., 2016; Kinseth et al., 2007). It is of note that release of IL-1ß is also dependent on GORASP proteins in LPS activated macrophages (Chiritoiu et al., 2019). GORASP is a key to unravelling this process because so far it is the only proteins that is required for many types of unconventional secretion of several different proteins in organisms through evolution, from yeast to mammals (add refs here).

In yeast, CUPS is marked by the presence of the single GORASP orthologue, Grh1 (Bruns et al., 2011). CUPS forms independent of COPI and COPII proteins, but requires the function of the phosphatidylinositol (PI) 4-kinase, Pik1, of the late Golgi membranes or trans-Golgi network (TGN) (Cruz-Garcia et al., 2014). During the time course of starvation, and correlating with the timing of unconventional secretion, CUPS “mature”, ultimately acquire large enveloping membranes in a process that requires the function of the PI 3-kinase, Vps34 and a subset of ESCRT proteins. In the absence of Vps34 or ESCRT complexes I, II, or III CUPS initially form, but later fragment. The major subunit of ESCRT-III, Snf7, also transiently localizes to the CUPS (Curwin et al., 2016).

We now report that that Drs2, a PI4P effector that functions as an aminophospholipid flippase and localized at the TGN in growth, is essential for CUPS biogenesis. This requirement in CUPS biogenesis is dependent on its binding partner Rcy1. Our data reveal that during unconventional secretion, cells create a new, TGN-derived compartment that is enriched in Drs2, Tlg2 (t-SNARE) and Snc2 (v-SNARE), which transiently contacts Grh1 containing CUPS. We have called this new compartment TCUPS for Touching CUPS. 4D imaging of cells by SCLIM revealed that CUPS and TCUPS make numerous, but transient contacts. Tubules emanating from TCUPS are often collared by CUPS. In some cases, the tubule appears to be severed post-contact. We believe these contacts are essential during the stress of starvation to facilitate compartment formation, maturation and stability of both CUPS and TCUPS. The discussion of our findings follows.

## Results

### PI4P effector Drs2 is necessary for CUPS biogenesis

PI4P is produced at the TGN by the PI 4-kinase Pik1 and is essential for proper Golgi function via various PI4P dependent pathways (Graham and Burd, 2011; Walch-Solimena and Novick, 1999). By use of a temperature sensitive allele, Pik1 was previously shown to be required for efficient CUPS biogenesis, however a PI4P fluorescent sensor does not localize to the Golgi, but is diffusely dispersed in the cytoplasm under these starvation conditions, indicating a decrease of Golgi PI4P levels (Cruz-Garcia et al., 2014). This is in accordance with published work indicating that glucose starvation leads to a rapid decrease of Golgi PI4P via re-localization of the enzymes, Pik1 and the Sac1 PI 4-phosphatase (Demmel et al., 2008; Faulhammer et al., 2007). This therefore begs the question: what is the function of the late Golgi and Pik1 in the overall process of CUPS formation and what are the effectors of PI4P in this pathway?

The multi-spanning transmembrane protein Drs2, aminophospholipid flippase, is a PI4P effector localized at the TGN membranes. We examined location of genomically expressed Grh1-2xmCherry and Drs2-3xGFP in growth and throughout the time course of starvation by confocal live spinning disk microscopy. In growth, Drs2 labelled 4-6 punctae per cell that were often apposed to but not colocalized with Grh1 (Figure 1A). Upon starvation, Grh1 re-localized to 1-3 larger foci, which we have shown previously to be the CUPS. Curiously, Drs2 also re-localized to 1-3 larger foci per cell and in addition displayed faint diffuse localization throughout the cytoplasm (Figure 1A). The foci of Grh1 and Drs2 were never observed to be stably localized, however, transient co-localization of the foci was frequently observed, particularly early in starvation (26% of cells). The rate of transient co-localization decreased throughout the time course of starvation (10% of cells), however this could be a reflection of the fact that the overall number of Drs2 compartments also decreased throughout starvation. The average number of Drs2 structures per cell decreased from 2.3/cell in the first 45 min to 1.5/cell in the last 30 min of starvation, while the average percentage of cells with no Drs2 structures increased from 7.9% to 13.8% in the same time period (Figure 1).

**Figure 1.**
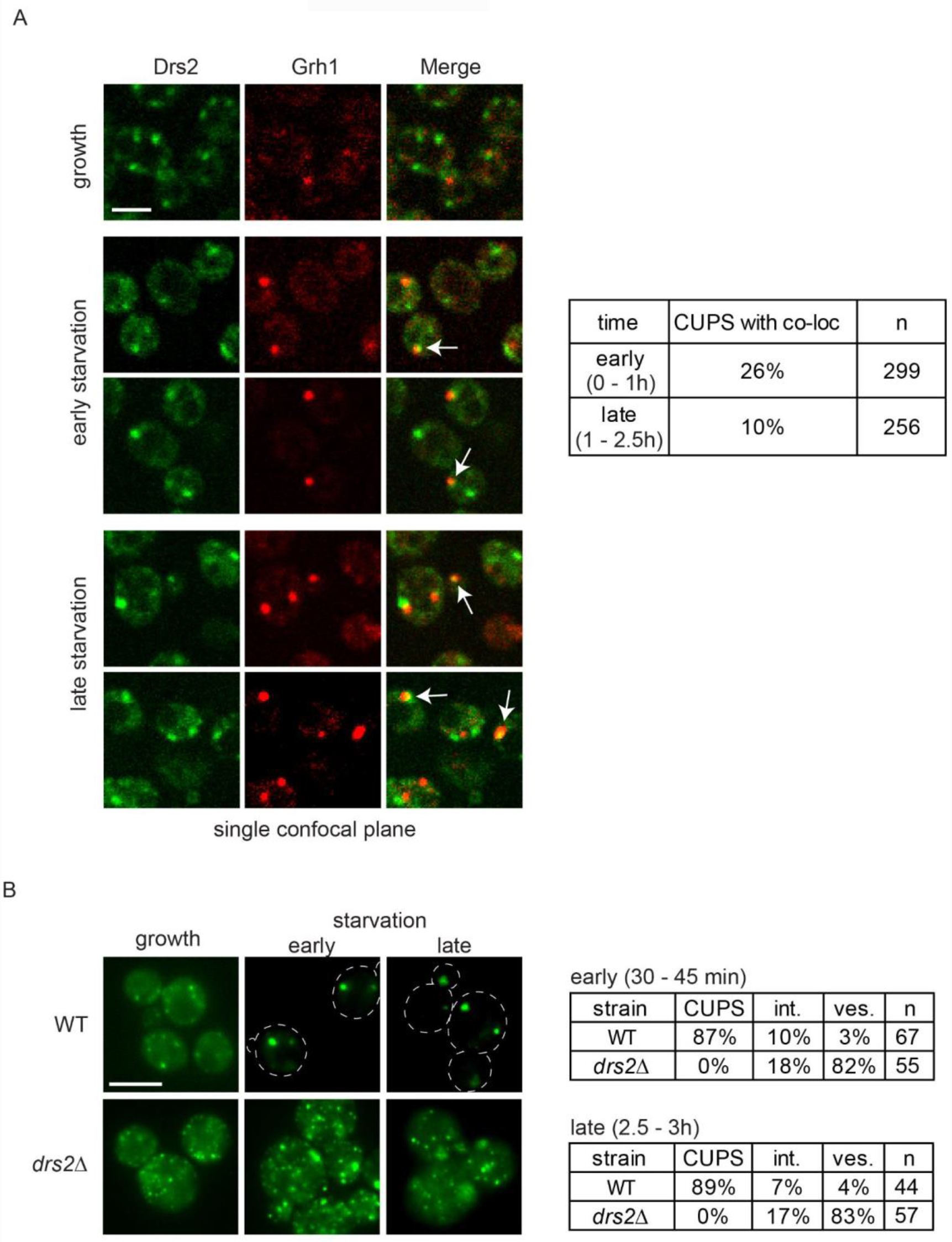
Drs2 is required for CUPS biogenesis. (A) Cells genomically expressing Drs2-3xGFP and Grh1-2xCherry were visualized by confocal spinning disk microscopy in growth conditions and starvation by incubation in 2% potassium acetate. Short movies were acquired at 10 second intervals to assess the frequency and duration of co-localization. Scale bar = 2μm. (B) Wild type and *drs2Δ* cells expressing Grh1-2xGFP were visualized by epifluorescence microscopy in growth conditions and after incubation in 2% potassium acetate for the indicated times. Cells were classified with normal CUPS (1-3 larger foci per cell); intermediate CUPS (“int.”), where a large focus is observed in addition to smaller structures; and vesiculated CUPS (“ves.”) where only small foci of Grh1 are observed. Scale bar = 2μm.

Next, we asked if Drs2 contributed to the process of CUPS formation. We examined localization of Grh1-2xGFP in cells lacking Drs2 and found a strong defect in CUPS biogenesis as observed by Grh1 localized to numerous smaller structures (Figure 1B). After 2.5-3 hours of starvation, CUPS were still unable to form in the absence of Drs2.

### Drs2 functions specifically with Rcy1 in CUPS formation

Drs2, a multi-spanning transmembrane protein functions at the TGN to flip mainly phosphatidylserine (PS), but also phosphatidylethanolamine (PE) to maintain phospholipid asymmetry and drive vesicle formation (Chen et al., 1999; Gall et al., 2002; Hua et al., 2002; Liu et al., 2008; Natarajan et al., 2004). The C-terminal domain of Drs2 has an autoinhibitory function that is relieved specifically upon binding of PI4P. Interaction of Drs2 with Arf-GEF Gea2 and Arf-like GTPase Arl1 are also critical in regulating multiple clathrin-dependent, anterograde pathways (Bai et al., 2019; Hankins et al., 2015; Natarajan et al., 2009; Timcenko et al., 2019; Tsai et al., 2013; Zhou et al., 2013). Clathrin and the PI4P sensor dissociate from the TGN upon starvation, we therefore did not expect a role for these players in CUPS formation.

Regardless, Gea2, Arl1 and clathrin (clathrin heavy chain or adaptor proteins) mutant strains lacking the function of these proteins were tested and revealed no effect on CUPS formation (Figure S1). Drs2, via its interaction to Rcy1, also regulates a retrograde pathway required for recycling of exocytic v-SNAREs Snc1 to the TGN. This interaction is also via the C-terminal domain of Drs2 in a region proximal to the PI4P binding site, and partially overlapping with the Gea2 binding site (Furuta et al., 2006; Hanamatsu et al., 2014). Deletion of *RCY1* resulted in highly vesiculated Grh1-positive structures, exactly as for loss of Drs2, clearly indicating the Drs2-Rcy1 branch of Drs2 function is specifically required for CUPS formation (Figure 2A).

**Figure 2.**
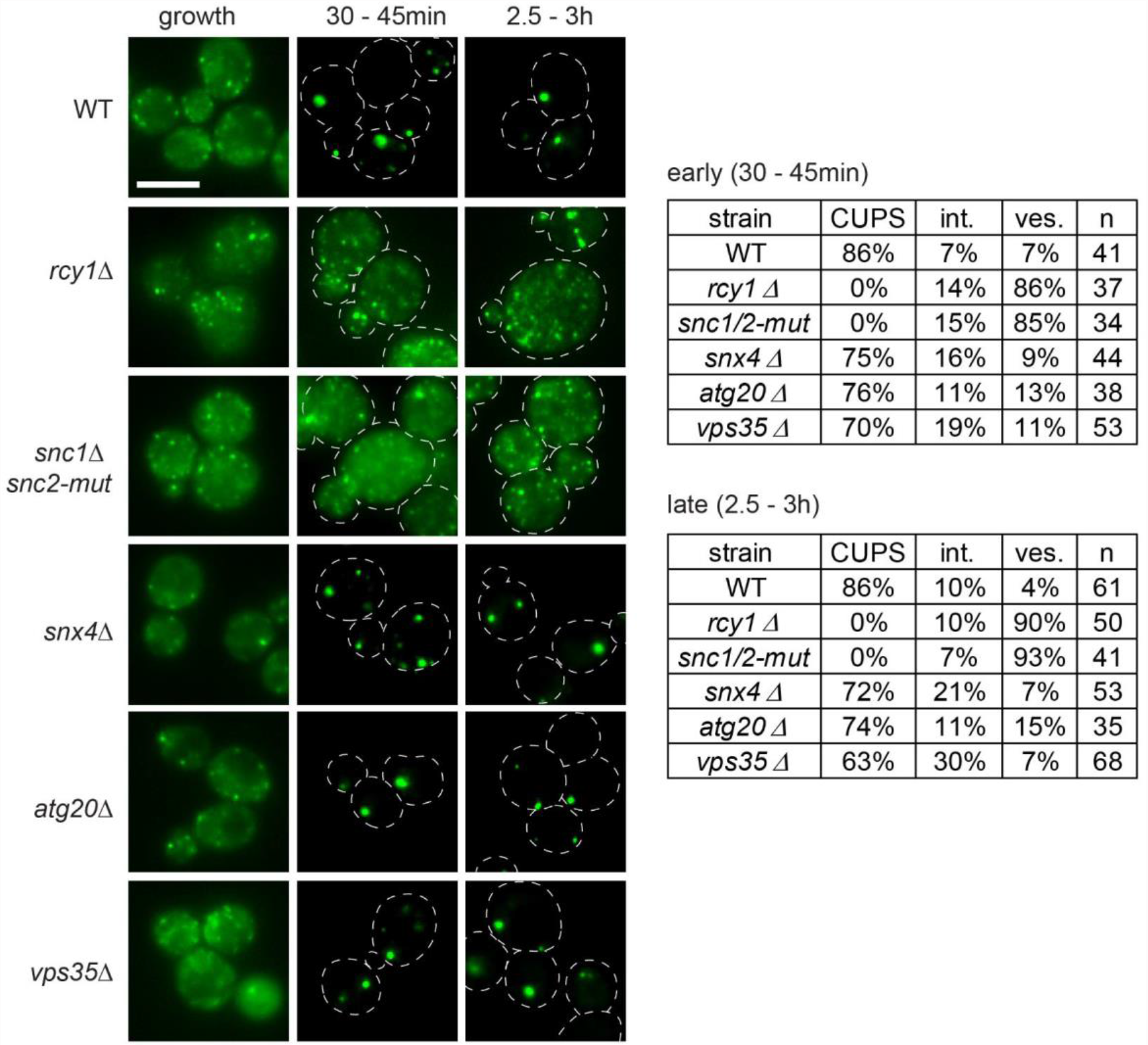
Drs2-Rcy1 pathway and the v-SNAREs, Snc1 and Snc2, are required for CUPS formation. Wild type and the indicated deletion or mutant strains expressing Grh1-2xGFP were visualized by epifluorescence microscopy in growth conditions and after incubation in 2% potassium acetate for the indicated times. Cells were classified with normal CUPS (1-3 larger foci per cell), intermediate CUPS (“int.”); where a large focus is observed in addition to smaller structures, and vesiculated CUPS (“ves.”); where only small foci of Grh1 are observed. Scale bar = 2μm.

### CUPS formation requires v-SNARE function

A major known function of the Drs2-Rcy1 pathway is recycling of the exocytic v-SNARE Snc1, we asked if Snc1 is also required for CUPS formation. Snc1 and Snc2 are the only post-Golgi v-SNARES in yeast and form an essential pair with redundant functions in cell growth and secretion (Protopopov et al., 1993). Single gene deletion of either produced no phenotype in CUPS formation (Figure S1). However, a double mutant temperature-sensitive strain lacking Snc1 and with Snc2 mutated to be inefficiently recycled from the plasma membrane, *snc2-V39A,M42A,* (Shen et al., 2013) displayed the highly vesiculated CUPS phenotype, even without temperature shift (Figure 2). The double mutant cells grew in normal conditions, but the sole mutated v-SNARE, Snc2, did not support formation of CUPS upon starvation.

Is the defect in CUPS due to a defect in recycling of v-SNAREs to the TGN or more directly related to the function of Drs2/Rcy1? Yeast have a minimal endomembrane system and an early TGN, marked specifically by the presence of the t-SNARE Tlg2, likely serves as an early endosome, while a later TGN is the site of exocytosis and clathrin-coated vesicle formation (Day et al., 2018; Tojima et al., 2019). The recycling of Snc1 to the early TGN has been shown to follow 3 distinct pathways; Drs2-Rcy1, that sort the Snc1 from the early endosome-like TGN, while the sorting nexins Snx4 and Atg20, as well as retromer, sort v-SNAREs at late endosomes (or the prevacuolar compartment in yeast), although retromer likely only becomes important when the other 2 pathways are not functioning (Best et al., 2020; Hanamatsu et al., 2014; Ma and Burd, 2019). We tested these pathways in CUPS biogenesis and observed no defect in cells lacking the sorting nexins Snx4 and Atg20, or the retromer subunit, Vps35 (Figure 2). Therefore, the defect observed in Drs2/Rcy1 deleted cells is not simply due to loss of v-SNARE pool at the TGN. The combined data suggest that CUPS specifically require membranes from an early endosome-like TGN compartment, in Drs2/Rcy1 dependent manner.

### Rcy1 and Snc1/Snc2 are required for unconventional secretion

Unconventional secretion in yeast leads to some secreted proteins being trapped in the cell wall or periplasmic space and to measure this secretion a mild cell wall extraction procedure is necessary to prevent cell lysis associated with perturbations to the rigidity of the cell wall (Curwin et al., 2016). As such, any genetic mutations or treatments (such as temperature shift) that exacerbate this problem of lysis cannot be tested by this assay to score unconventional secretion. Cells lacking Drs2 have numerous defects, particularly in lipid homeostasis (Hankins et al., 2015) and therefore could not be tested (data not shown). However, *rcy1Δ* cells do not exhibit as many defects associated with loss of Drs2 function, therefore we tested their capacity to secrete unconventional cargoes such as Acb1, and the antioxidants Sod1 and Trx2. Wild type and *rcy1Δ* cells were starved for 2.5 hours after which the secreted material was extracted from the cell wall, as described previously (Curwin et al., 2016). The intracellular and secreted fractions were probed by western blot for the various cargoes, Cof1-which is used to monitor cell lysis, and the known cell wall protein Bgl2. Loss of Rcy1 led to a strong defect in release of Acb1, Sod1 and Trx2 without causing release of cytoplasmic content measured by the lack of Cof1 presence (Figure 3A). Similarly, the v-SNARE double mutant defective in CUPS formation (Figure 2) was tested and also exhibited a reduction in secretion of Acb1, Sod1 and Trx2 (50-60% compared to control cells) (Figure 3B). Therefore, we can conclude that unconventional secretion in starvation requires Rcy1 (presumably in concert with Drs2) and v-SNARE activity, but the precise function of these players remains to be determined.

**Figure 3.**
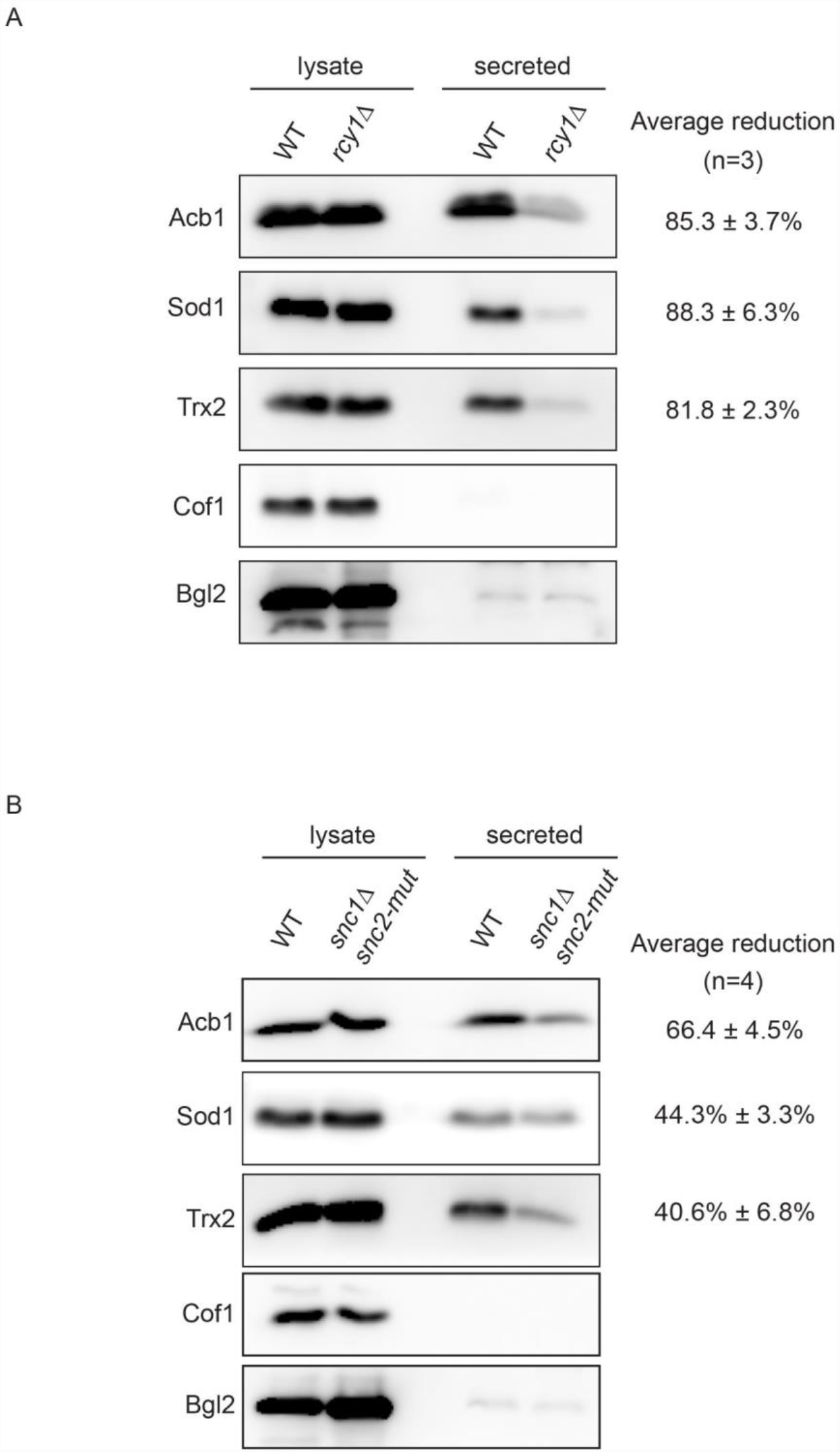
Rcy1 and v-SNAREs are required for unconventional secretion. (A+B) Wild type, *rcy1Δ* or *snc1Δ snc2-V39A, M42A* cells were grown logarithmic phase, washed twice and cultured in 2% potassium acetate for 2.5 hours. The cell wall proteins were extracted from equal number of cells followed by precipitation with TCA (“secreted”). Lysates and secreted proteins were analyzed by western blot and the ratio of the secreted/lysate for the indicated protein was determined and compared to that of wild type in each experiment. Statistical analyses were performed for the indicated unconventional cargo proteins and the reduction in secretion compared to wild type is indicated ± standard deviation.

### Drs2 and the SNARE proteins Snc2 and Tlg2 label a new compartment that transiently contacts CUPS

We generated N-terminal GFP fusions of the v-SNAREs, Snc1 and Snc2, and their cognate t-SNARE, Tlg2, which preferentially labels the early TGN and therefore likely receives the v-SNARE vesicles being recycled in a Drs2-Rcy1 dependent manner. Most analyses of v-SNARE itinerary have been performed by overexpression of an N-terminal GFP tagged Snc1, that at steady-state labels mostly the plasma membrane of growing buds and some internal structures (Lewis et al., 2000). Recently, Graham and colleagues generated an mNG-Snc1 construct expressed at much lower levels, which preferentially labelled the TGN and endosomes (Best et al., 2020). To avoid plasmid overexpression altogether, we integrated the GFP tag at the N-terminus of each SNARE, at its endogenous locus, under the control of Sed5 promoter. Interestingly, a different steady state pattern for Snc1 and Snc2 was observed during growth. The overall signal of Snc1 was weaker than that of Snc2, and distinct localization of Snc1 was only observed in the tips of very small budded cells and somewhat in the necks of large budded cells (Figure 4A). Unbudded and large/medium budded cells displayed mostly a diffuse signal of Snc1. Snc2, on the other hand, exhibited distinct localization in all cells, preferentially labelling mostly internal structures, the neck of large budded cells, and occasionally the plasma membrane of small and medium budded cells (Figure 4A). So, although the v-SNAREs are redundant in function they clearly have their own preferred steady-state itineraries. Tlg2 localized as expected in growth, labelling 4-6 punctae per cell (Figure 4A). None of the SNARE proteins could be co-localized with Grh1 in growth conditions.

**Figure 4.**
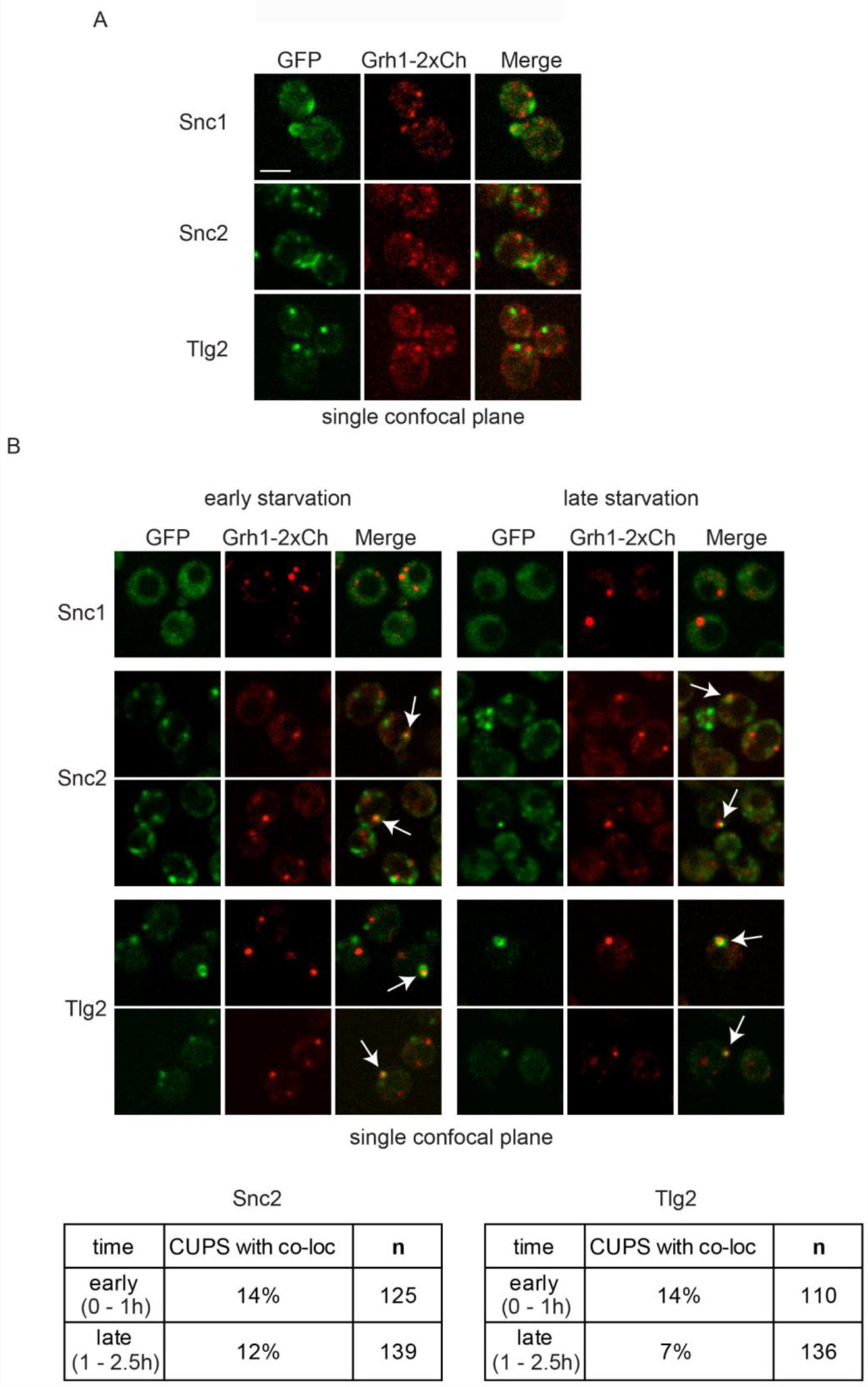
CUPS contain a pool of the v-SNARE Snc2 and the t-SNARE Tlg2. (A) Cells genomically expressing Grh1-2xCherry with GFP-Snc1, GFP-Snc2 or GFP-Tlg2 were visualyzed by confocal spinning disk microscopy in growth conditions and (B) throughout the time course of culture in 2% potassium acetate. Short movies were acquired at 10 second intervals to assess the frequency and duration of co-localization. Scale bar = 2μm.

Upon starvation the Snc1 signal rapidly became diffuse in most cells (less than 5% retained 1-2 faint foci) indicating Snc1 is the preferred v-SNARE for exocytosis. In contrast, the Snc2 signal remained high in all cells, labelling fewer and larger punctate elements (Figure 4B). Tlg2 also labelled fewer and larger structures immediately upon starvation, in the same manner as Drs2. Both Snc2 and Tlg2 structures could be found transiently co-localized with Grh1 (Figure 4B). In the case of Snc2, this was observed on average in 13% of cells at any particular time point in starvation, whileTlg2 co-localization with Grh1 was more frequent early in starvation (14% of cells early and 7% of cells later in starvation), similar to Drs2. Examination of GFP-Tlg2 with Drs2-3xCherry revealed that they are indeed contained in the same compartment in starvation (Figure S2). Drs2 could be predicted to be in both the early and late TGN membranes due its function in anterograde and retrograde transport, while Tlg2 is specific to early TGN. We observed a partial co-localization in growth conditions, as expected if this were true (Figure S2). In starvation, the number of TGN membranes was reduced and the co-localization of Drs2 and Tlg2 was greatly increased (Figure S2). The same was also observed when mCherry-Snc2 and GFP-Tlg2 were tested for their location during starvation. The signal of mCherry-Snc2 was very weak compared to the GFP version, but larger Snc2 structures were observed to co-localize with Tlg2 (Figure S2). Therefore, starvation induces the formation of a new compartment derived from the early TGN that is enriched Drs2, Tlg2 and Snc2. We now discuss data showing this new compartment contacts CUPS transiently and based on this feature we have called it TCUPS for Touching CUPS.

### SCLIM reveals the process of CUPS formation in 4D

We previously presented the ultra-structure of CUPS using CLEM (correlative light electron microscopy) as a spherical tubulovesicular structure that grows in overall size during starvation and acquires a large enveloping cup-shaped cisternae (Curwin et al., 2016). To gain better insight into the organization of membranes that compose CUPS and its potential interaction with TCUPS we used super-resolution confocal live imaging (SCLIM) (Kurokawa et al., 2013, 2019). SCLIM analysis of Grh1-2xGFP has confirmed these structures and further revealed their dynamic behaviour (Figure 5 and Movies1-4). Grh1 was localized to many small and mobile structures in growth conditions. Detailed analysis of larger Grh1-positive structures at 3 hours starvation revealed mature CUPS could be categorized in 3 forms; spherical, cup-like and curved (Figure 5B). Even though the overall mobility of structures decreased during the time course of starvation, the CUPS morphology could dynamically change between the different forms (Movie 1). Dynamic Grh1 structures could be observed to contact each other at times, possibly fusing and becoming more stable (Figure 5C and Movies1-4). Moreover, a subsequent analysis earlier in starvation (1 - 1.5 hour), indicated dynamic Grh1-positive structures contacted numerous times, growing in size likely by fusion, as well larger Grh1 structures could also be observed to fragment at times (Movies S1-3). Altogether, the SCLIM analyses reveal that CUPS form by dynamic interactions between Grh1 containing membranes, which likely involves fusion and fission. These highly dynamic interactions then reach a steady state and generate a more stable CUPS.

**Figure 5.**
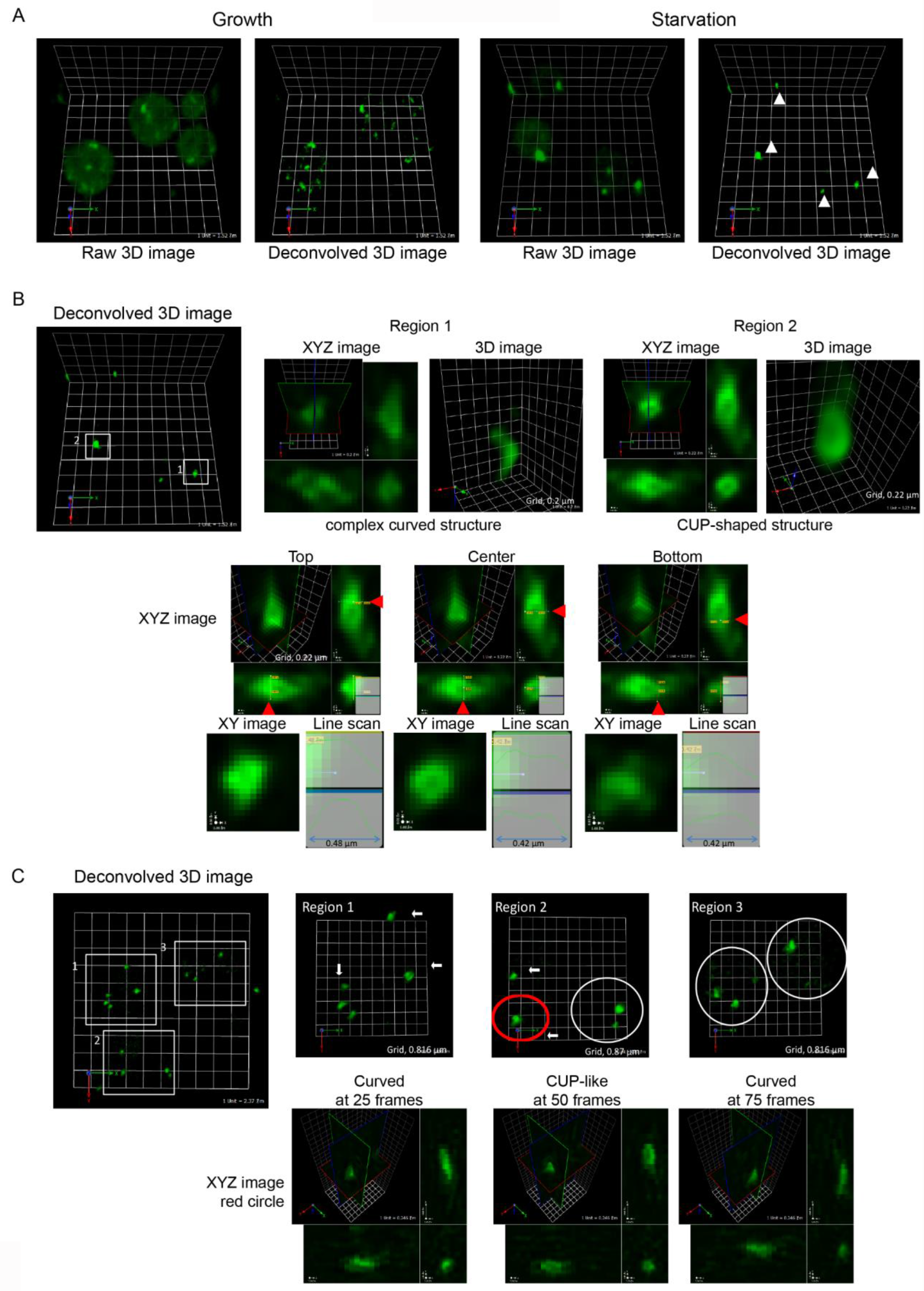
SCLIM reveals the dynamic structure of CUPS. (A) Grh1-2xGFP cells were incubated in normal growth conditions or 2% potassium acetate for 3 hours and visualized by SCLIM. In growth, Grh1 labelled many small and mobile structures (early Golgi membranes and ER-exit sites). In starvation, Grh1 labelled fewer, larger, and less mobile membrane structures (CUPS) Grid = 1.52μm (B) Line scan analysis in 3D of multiple CUPS structures revealed three forms; spherical (3/14), complex curved (8/14) or cup-shaped (3/14). (C) Visualization of CUPS over time showed stable, mature CUPS are still dynamic, able to change morphology between the different forms. Arrows = non-moving structures; dotted circles = moving structures. Red circle is the region in the time-lapse images at 25, 50 and 75 frames. Movies 1-4; regions 1-3 and red circle region. 20 sec intervals, 60x speed.

### Dynamics of Drs2 compartment (TCUPS) and contacts to CUPS

Before monitoring Grh1 in combination with other proteins by SCLIM, we observed the changes in the TGN compartments induced upon starvation and the formation of TCUPS. We therefore examined of the organization of Drs2-3xmCherry and GFP-Tlg2 in growth and starvation. Tlg2 localizes to an early TGN compartment while. In growth, co-localization of Drs2 and Tlg2 was observed in some cisternae, but not all. Following the structures over time revealed an order of events, with structures first labelled by Tlg2, then acquiring Drs2 (co-localization), subsequently losing Tlg2, and finally the loss of Drs2 (Figure 6A and Movie 5). In starvation, the extent of Drs2 and Tlg2 co-localization was greatly increased as TCUPS form. Interestingly, the same dynamics could be observed in starvation, with structures only marked by Tlg2 then acquiring Drs2, followed by Drs2 alone, and finally the loss if TCUPS compartment (Figure 6B and Movies 6-8).

**Figure 6.**
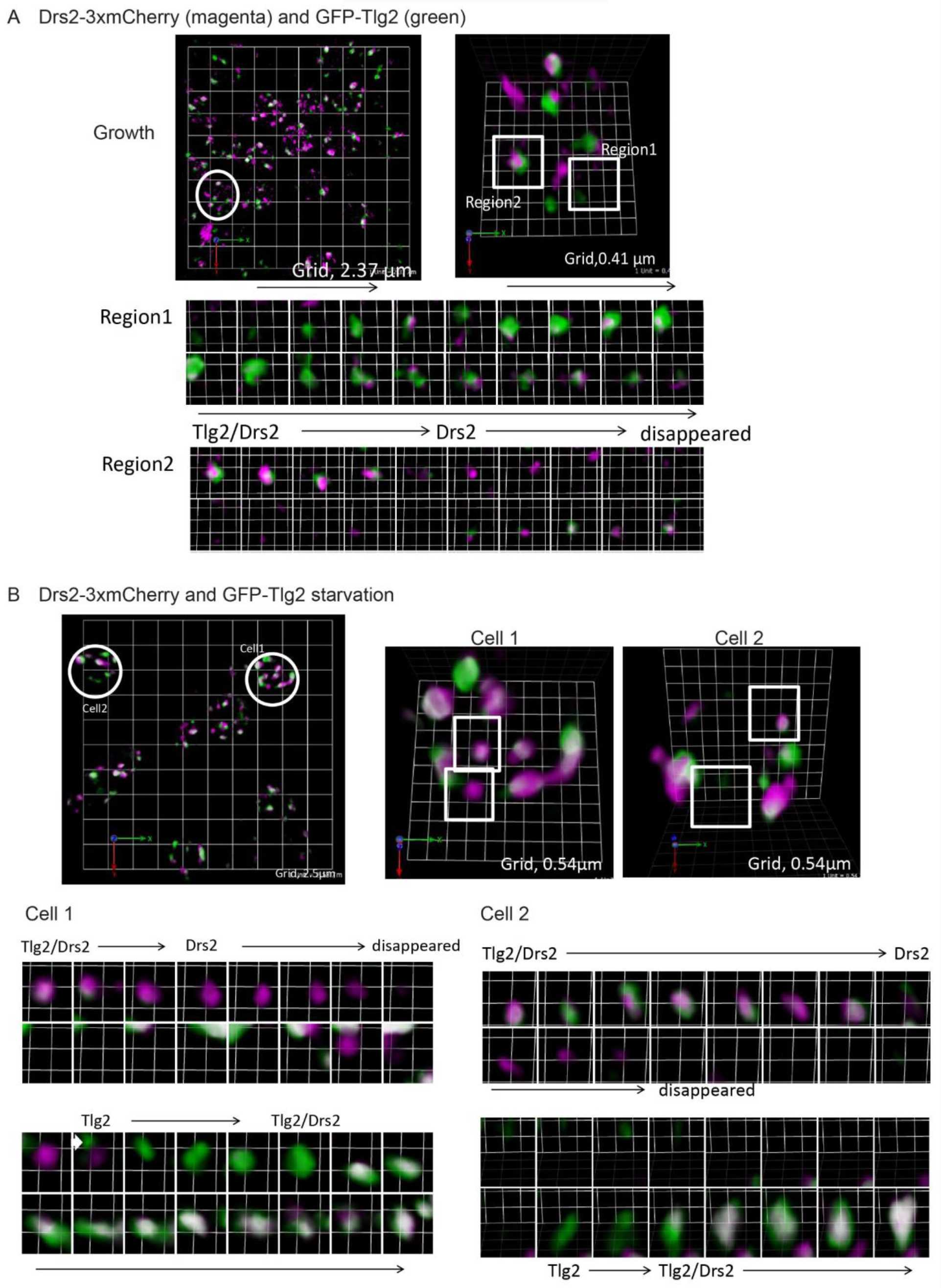
SCLIM analysis of Drs2 and Tlg2 labelled structures in growth (TGN) and starvation (TCUPS) (A) Drs2-3xCherry (magenta) and GFP-Tlg2 (green) cells were visualized in growth condition. White indicates co-localization. Time-lapse images of the two regions indicated. Movie 5, 4 sec intervals, 20x speed. (B) Drs2-3xCherry (magenta) and GFP-Tlg2 (green) cells were visualized at 2 hours starvation. White indicates co-localization. Time-lapse images of the two cells indicated. Movie 7-zoomed out and Movies 8-9 – zoomed in of two indicated regions, 10 sec intervals, 20x speed.

### Fragmentation of TCUPS by contact with CUPS

Finally, we analyzed the dynamics of Grh1-2xmCherry containing CUPS in combination with Drs2-3xGFP, GFP-Tlg2 or GFP-Snc2 to visualize the CUPS-TCUPS interaction. SCLIM analysis captured numerous contacts between CUPS and TCUPS. These contacts were not observed frequently, but given they are transient they are likely difficult to capture. The nature of contacts often involved insertion of a TCUPS tubule into CUPS. This could be observed with Drs2, Tlg2 or Snc2 as the marker of TCUPS. In Figure 7A and Movie 9 an example with Drs2 is provided, where the Drs2-positive membranes inserted into CUPS and fragments were produced. In another remarkable movie, with Tlg2 as the TCUPS marker, the CUPS collar and appear to sever a tubule derived from TCUPS, although direct cutting of TCUPS by CUPS cannot be conclusively stated by this analysis alone (Figure 7B and Movie 10). In addition to TCUPS, Snc2, and to a lesser extent Drs2, also labelled numerous smaller structures that also often contacted with or were in the vicinity of CUPS (Figure S3 and Movies S4-8). These are likely vesicles/tubules, as Drs2 itself has been shown to be packaged into vesicles during its activity (Liu et al., 2008). The combined evidence suggests that TCUPS is being consumed during starvation in the process of vesicle formation, driven, at least partially, by the action of Drs2-Rcy1. Although these vesicles are not fusing directly to the CUPS, as we do not observe this by SCLIM, the absence of Drs2-Rcy1 activity is essential for both CUPS formation and unconventional secretion.

**Figure 7.**
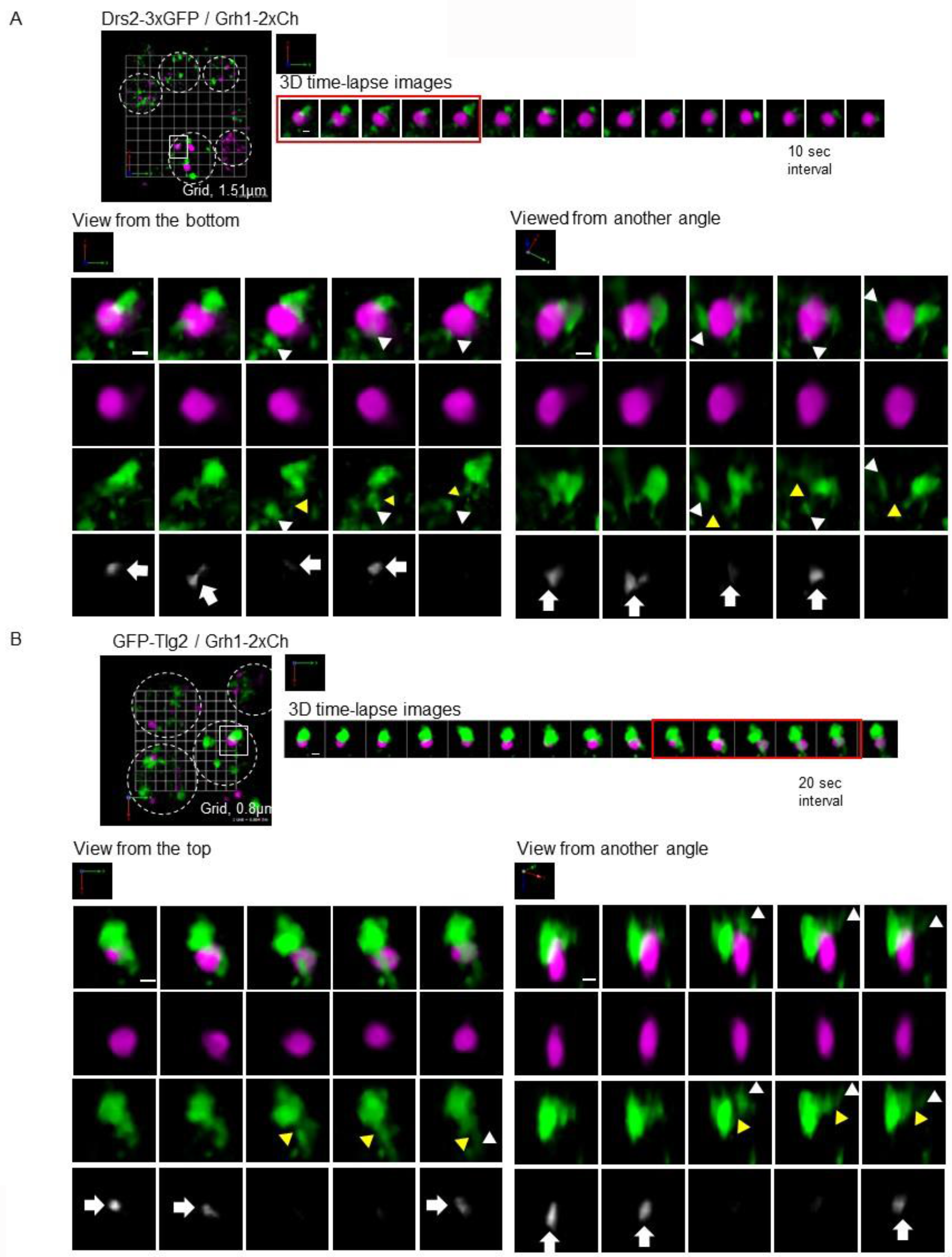
SCLIM analysis of CUPS-TCUPS contacts. (A) Grh1-2xCherry (magenta) and GFP-Tlg2 (green) cells cultured in starvation condition 1.5 hours. 3D time-lapse images (20 sec intervals). White arrowheads show separated membrane structures labeled with GFP-Tlg2. Yellow arrowheads indicate where the membrane structures have been cut. White arrows indicate where Grh1 contacts with Tlg2 protrusive membrane. Scale bar=0.5µm. Movie 9, 20 sec intervals, 20x speed. (B) Grh1-2xCherry (magenta) and Drs2-3xGFP (green) cells cultured in starvation condition 1.5 hours. 3D time-lapse images (10 sec intervals). White arrowheads show separated membrane structures labeled with Drs2-3xGFP. Yellow arrowheads indicate where the membrane structures have been cut. White arrows indicate where Grh1 contacts with Drs2 protrusive membrane. Scale bar=0.5µm. Movie 10, 10 sec intervals, 20x speed.

## Discussion

George Palade mapped the pathway of protein secretion, which laid the foundation for decades of research on how proteins are exported from the ER and transported via the Golgi complex to their ultimate destinations. Blobel revealed how proteins enter this ER-Golgi pathway of secretion via the N-terminal signal sequence. The knowledge that eukaryotic cells secrete proteins lacking the N-terminal signal sequence and therefore cannot follow the Blobel-Palade’s pathway of protein secretion is also beginning to receive attention. This unconventional secretory pathway releases a large and diverse class of proteins with varied vital physiological functions in extracellular space for immune surveillance, tissue reorganization, insulin homeostasis, and protection from oxidative damage, for example. We showed that the Golgi associated protein GORASP, is required for the secretion of Acb1 in nutrient-starved *Dictyostelium discoideum* (Kinseth et al., 2007). This function of GORASP (a single gene in invertebrates, two genes in vertebrates) is now known to be conserved. We have focused on the pathway of GORASP-dependent unconventional secretion and our findings have led to identification of a compartment that we call CUPS, which is synthesized under the conditions that trigger unconventional secretion (Bruns et al., 2011; Cruz-Garcia et al., 2014). CUPS are composed of membranes recruited from the Golgi apparatus, identified by the presence of Grh1 (GORASP ortholog in yeast), and form without the function of COPI and COPII components. CUPS formation depends on the phosphoinositides PI4P for biogenesis and PI3P for its stability. The major ESCRT-III protein Snf7 is also found at the CUPS transiently (Curwin et al., 2016). Importantly, all of these molecular components are also required for secretion of Acb1, Sod1 and Trx2 (Cruz-Garcia et al., 2020; Curwin et al., 2016).

Our new findings reveal that PI4P functions in the requirement of Drs2, a transmembrane aminophospholipid flippase, in both CUPS formation and unconventional secretion. Of the many proteins that function with Drs2 in trafficking at the TGN that includes Gea2, Arl1 and Rcy1, only Rcy1 is involved in unconventional secretion. We also report a requirement for v-SNARE function (Snc1 and Snc2 orthologous pair) in CUPS formation and unconventional secretion. Together, the starving yeast generate a single new compartment from the TGN that contains Drs2, the v-SNARE Snc2 (but not Snc1) and the t-SNARE Tlg2. This compartment that we have called TCUPS contacts Grh1 containing CUPS.

### Building and remodeling TCUPS during unconventional protein secretion

Live cell 4D imaging (SCLIM) has revealed membrane contacts between CUPS and TCUPS. The contacts are transient and highly dynamic, with CUPS membranes often observed to enwrap or encircle TCUPS. We have captured a fascinating event in the contact of CUPS to TCUPS: a tubule emerging from TCUPS is collared by CUPS and appears to be severed. This event is reminiscent of the contact of ER and the endosomes and the fission of the latter compartment (Rowland et al., 2014). We have previously shown that Snf7, a protein of the ESCRT III complex, is recruited transiently to CUPS. However, Vps4 is not required for CUPS biogenesis or for the secretion of Acb1. We propose that CUPS collar the neck of a TCUPS derived tubule and Snf7 located at this junction is involved in events leading to the severing of the respective tubule. There is at least one example of ESCRT mediated severing of a tubule on the cytoplasmic face of a membrane bound compartment (McCullough et al., 2015).

### Unconventional secretion aping conventional secretion

In Palade’s conventional secretory pathway, proteins destined for secretion are translocated into the ER lumen, folded and sorted, transported to the Golgi, and from there on to their final destinations. Unconventionally secreted cargoes are not glycosylated so they do not need to enter the ER. But there is still the question of how a fraction of this class of proteins is selected and translocated from the cytoplasm for their subsequent delivery to the extracellular space. Our data reveal that membranes from the Golgi and the TGN are extracted by COPI-, COPII-, and clathrin-independent pathway to create two distinct compartments (Figure 8). These are marked by: 1, Grh1 (previously defined as CUPS) and 2, Drs2/Tlg2/Snc2 (TCUPS). CUPS and TCUPS are therefore the early and late Golgi equivalent of the unconventional secretory pathway. The only obvious difference being that they do not contain the Golgi specific glycosylating enzymes.

**Figure 8.**
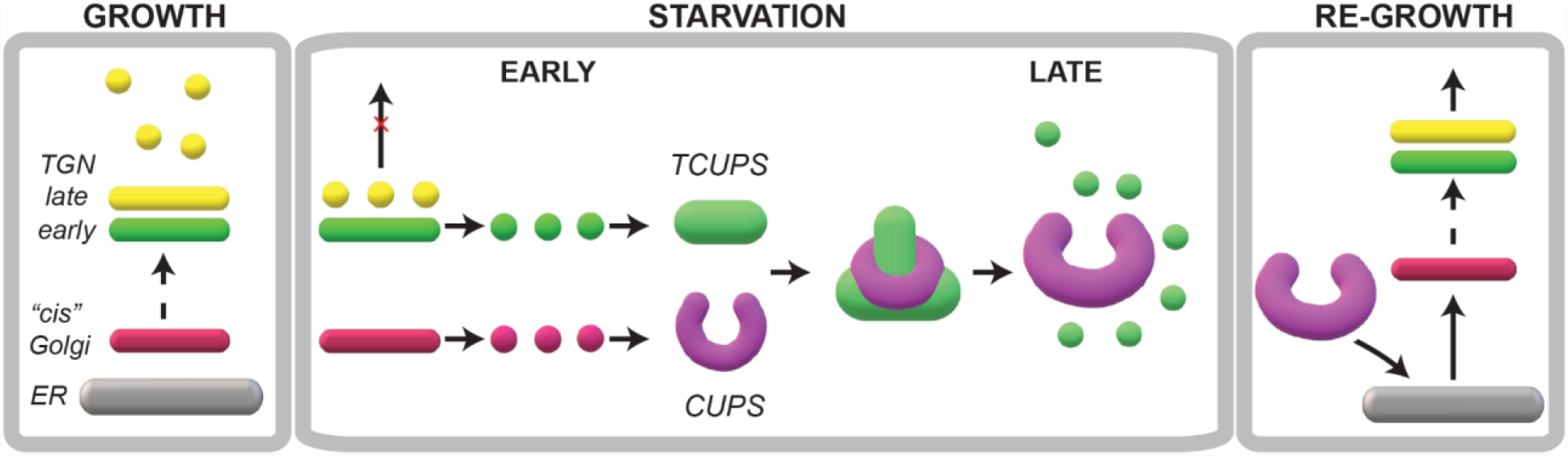
A working scheme building CUPS-TCUPS for unconventional secretion. During growth, cells predominantly depend on the conventional ER-Golgi pathway of protein secretion. When cells are cultured in starvation medium, there is a sharp reduction in the use of conventional secretory pathway and the cells switch to a new or an unconventional mode to release essential proteins to cell’s exterior. A cis Golgi membrane produces small fragments, that do not contain glycosylation enzymes, in a COPI independent manner to synthesize CUPS (magenta). The early TGN produces small membranes to generate a compartment that we have called TCUPS (green). Our data show that tubules emanating from TCUPS are collared by CUPS, which is followed by severing of the tubule. We suggest that these contacts, over a period, lead to the consumption of TCUPS to produce smaller elements (vesicles + tubules). These smaller elements are likely used for delivering essential proteins to other compartments of the cell and release proteins like SOD1 and Acb1 to cell’s exterior. This mode of TGN consumption is common to both the conventional and unconventional protein secretion processes. Upon shifting cells to growing conditions, components of the CUPS are delivered by COPI vesicles to the ER, which then traffic the respective components to the Golgi, thereby restoring the Golgi to restart the conventional mode of protein secretion.

We have shown previously that CUPS contain Acb1, but this has only been observed with immunoelectron microscopy. We admit that we do not know if Acb1 enters directly into CUPS. CUPS and TCUPS transiently contact and it is possible that Acb1 is transferred from CUPS to TCUPS during their transient connection. TCUPS, we suggest are the sorting station for the cargo. How this is achieved is also not known. We have seen that CUPS collar a tubule emanating from TCUPS. Snf7 of the ESCRT pathway that is also recruited transiently to CUPS might be involved in generating and or severing this tubule from TCUPS. Are the tubules enriched in cargo destined to the cell surface? Over time, TCUPS are completely fragmented Much like the TGN during the conventional protein secretion. In the end this arrangement of creating new compartments is used by cells to deliver essential activities to the extracellular space during starvation. Upon return to the normal growth conditions, the remaining CUPS are reabsorbed to the ER by a COPI dependent manner. This is reminiscent of COPI dependent retrograde transport to the ER under growth conditions.

In sum, during starvation, the cells use the preexisting Golgi membrane to create new compartments for capturing cytoplasmic proteins, their sorting and the export for secretion. This foundational scheme provides a means to address the mechanism of unconventional protein secretion.

## Materials and Methods

### Yeast strains and media

Yeast cells were grown in synthetic complete (SC) media (0.67% yeast nitrogen base without amino acids, 2% glucose supplemented with amino acid drop-out mix from ForMedium). All strains are derived from the BY4741 background (*MATa his3Δ1 leu2Δ0 met15Δ0 ura3Δ0*). Deletion strains were from the EUROSCARF collection with individual genes replaced by KanMx4. Strains expressing C-terminally 2xyeGFP- and/or 2xyomCherry-tagged Grh1 were constructed by a PCR-based targeted homologous recombination and have been described previously (Cruz-Garcia et al., 2014). In many cases, strains were generated by mating and sporulation, followed by selection of clones with appropriate markers, and confirmation of haploidy. The double mutant v-SNARE (BY4741 *snc1Δ::kanMX4 snc2-V39A,M42A*) strain was provided by Peter Novick. This mutant expressing Grh1-2xyeGFP was generated by mating, sporulation and confirmation of markers and temperature sensitivity. Drs2-3xGFP, KanMx4 strain was a gift from Oriol Gallego (UPF).

#### Construction of N-terminally tagged SNAREs

The plasmid pYM-N9 (PCR toolbox) was used to generate a new template vector for PCR-based integration containing the NatNT2 selection cassette, the promoter of Sed5, followed by 2 tandem yeGFP. The promoter of Sed5 was amplified from genomic DNA with primers “SacI PrSed5 Fw1”: ATAGAGCTCTTACCATGTCCTCCAGAATTACGA and “XbaI PrSed5 Rv1”: TCATCTAGAGGGAGTTGTGTGGTATGGTG to generate a 658 bp fragment and was cloned into pYM-N9, replacing the high expression *ADH1* promoter. Subsequently a second yeGFP fragment was generated using primers “XbaI ATG yeGFP”: TGATCTAGAAAAAATGTCTAAAGGTGAAGAATTATTCACTGG and “EcoRV non-stop yeGFP”: TCTGATATCAGGCCTCATCGATGAATTCTCTGTCGGA and cloned downstream of the first yeGFP. Finally, standard S1/S4 primers were used to generate the N-terminal integration fragment targeting the Snc1, Snc2 and Tlg2 loci. Strains were confirmed as positive by microscopy and PCR to confirm the presence of 2 yeGFP. In most cases, however only one GFP integrated and the resulting 1xyeGFP strains were used. In the case of Snc1, 2xyeGFP was initially analyzed and subsequently a single yeGFP version was generated and found to behave in an identical manner as the 2xyeGFP version. SnapGene software (from GSL Biotech, Chicago, IL; available at www.snapgene.com) was used for molecular cloning design.

### Antibodies

All antibodies were raised in rabbit and have been described previously. Anti-Sod1 and anti-Trx2 were the kind of gifts of Yoshiharu Inoue (Research Institute for Food Science, Kyoto University) and T. O’Halloran (Northwestern University, Chicago, IL), respectively. Anti-Cof1 was kindly provided by John Cooper (Washington University in St. Louis) and anti-Bgl2 was a gift from Randy Schekman (UC Berkeley). Anti-Acb1 antibody was generated by inoculating rabbits with recombinant, untagged Acb1, purified from bacteria and has been described previously (Curwin et.al 2016). HRP conjugated anti-rabbit secondary was from Jackson Immunoresearch (Cat# 711-035-152).

### Cell wall extraction assay

Yeast cells were inoculated at a density of 0.003-0.006 OD_600_/mL in SC medium at 25°C. The following day, when cells had reached OD_600_ of 0.4-0.7 equal numbers of cells (16 OD_600_ units) were harvested, washed twice in sterile water, resuspended in 1.6 mL of 2% potassium acetate and incubated for 2.5 hours. When growing cells were to be analyzed 16 OD_600_ units were directly harvested. The cell wall extraction buffer (100mM Tris-HCl, pH 9.4, 2% sorbitol) was always prepared fresh before use and kept on ice. To ensure no loss of cells and to avoid cell contamination in the extracted buffer, 2mL tubes were siliconized (Sigmacote) prior to collection. Cells were harvested by centrifugation at 3000xg for 3 minutes at 4°C, medium or potassium acetate was removed and 1.6 mL of cold extraction buffer was added. Cells were resuspended gently by inversion and incubated on ice for 10 minutes, after which they were centrifuged as before, 3000xg for 3 minutes at 4°C, and 1.3 mL of extraction buffer was removed to ensure no cell contamination. The remaining buffer was removed and the cells were resuspended in 0.8 mL of cold TE buffer (Tris-HCl, pH 7.5, EDTA) with protease inhibitors (aprotinin, pepstatin, leupeptin (Sigma)) and 10 μL was boiled directly in 90 μL of 2x sample buffer (lysate). For western blotting analysis, 30 μg of BSA (bovine serum albumin (Sigma)) carrier protein and 0.2 mL of 100% Trichloroacetic acid (Sigma) was added to the extracted protein fraction. Proteins were precipitated on ice for 1 hour, centrifuged 16,000xg for 30 minutes and boiled in 50 μL 2x sample buffer. For detection, proteins (10 μL each of lysate or wall fractions) were separated in a 12% polyacrylamide gel before transfer to 0.2 μm nitrocellulose (GE Healthcare) for detection by western blotting. For preparation of cell wall extracts for mass spectrometry analysis, no BSA carrier protein was added and the proteins were precipitated with acetone and not TCA.

### Epifluorescence microscopy

After incubation in the appropriate medium cells were harvested by centrifugation at 3,000 g for 3 min, resuspended in a small volume of the corresponding medium, spotted on a microscopy slide, and imaged live with a DMI6000 B microscope (Leica) equipped with a DFC 360FX camera (Leica) using an HCX Plan Apochromat 100x 1.4 NA objective. Images were acquired using LAS AF software (Leica) and processing was performed with ImageJ 1.47n software.

### Spinning disk confocal fluorescence microscopy

After incubation in starvation medium for 20 min, ∼0.05 OD600nm of cells were plated in starvation medium on Concanavalin A–coated (Sigma-Aldrich) Lab-Tek chambers (Thermo Fisher Scientific) and were allowed to settle for 20 min at 25°C. Cells were continuously imaged up to 10 min throughout starvation. Whole cell Z stacks with a step size of 0.4 μm were continuously acquired (10 sec frames) using a spinning-disk confocal microscope (Revolution XD; Andor Technology) with a Plan Apochromat 100× 1.45 NA objective lens equipped with a dual-mode electron-modifying charge-coupled device camera (iXon 897 E; Andor Technology) and controlled by the iQ Live Cell Imaging software (Andor Technology). Some later images were taken on a newer spinning disk system (Andor Dragonfly) equipped with a 488nm and/or 561nm diode, using a U Plan Apo 60x 1.4 oil objective and an iXON-EMCCD Du-897 camera. A 2x camera zoom was used to reach Nyquist sampling, Fusion software was used for acquisition. Post-acquisition processing was performed with ImageJ 1.47n software.

### SCLIM (super resolution confocal live imaging)

For high-speed live imaging, yeast cells were immobilized on glass slides using concanavalin A and imaged by SCLIM. SCLIM was developed by combining Olympus model IX-71 inverted fluorescence microscope with a UPlanSApo 100× NA 1.4 oil objective lens (Olympus), a high-speed and high signal-to-noise ratio spinning-disk confocal scanner (Yokogawa Electric), a custom-made spectroscopic unit, image intensifiers (Hamamatsu Photonics) equipped with a custom-made cooling system, magnification lens system for giving 266.7× final magnification, and EM-CCD cameras (Hamamatsu Photonics) (Kurokawa et al., 2013). Image acquisition was executed by custom-made software (Yokogawa Electric). For 3D images, we collected optical sections spaced 100 nm apart in stacks by oscillating the objective lens vertically with a custom-made piezo actuator. Z stack images were converted to 3D voxel data and processed by deconvolution with Volocity software (Perkin Elmer) using the theoretical point-spread function for spinning-disk confocal microscopy. Imaging analysis was done using Volocity and MetaMorph software (Molecular Devices).

## Acknowledgements

We thank members of the Malhotra lab for their valuable discussions. V. Malhotra is an Institució Catalana de Recerca i Estudis Avançats professor at the Centre for Genomic Regulation. This work was funded by grants from the Spanish Ministry of Economy and Competitiveness (BFU2013-44188-P and BFU2016_75372-P to VM) and by Grants-in-Aid for Scientific Research from Japan Society for the Promotion of Science (JP17H06420, and JP18H05275 to AN and KK). We acknowledge support of the Spanish Ministry of Economy, Industry and Competitiveness (MEIC) to the EMBL partnership, the Programmes “Centro de Excelencia Severo Ochoa 2013-2017” (SEV-2012-0208 & SEV-2013-0347), and the CERCA Programme/Generalitat de Catalunya. This work reflects only the authors’ views, and the EU Community is not liable for any use that may be made of the information contained therein. The authors declare no competing financial interests.

## Author Contributions

Following the CRediT nomenclature, Amy J. Curwin, Nathalie Brouwers and Kazuo Kurokawa contributed to investigation, and visualization. AJC also contributed to conceptualization, formal analysis, project administration, methodology, validation, writing – original draft and writing – review & editing. Vivek Malhotra and Akihiko Nakano contributed to the conceptualization, funding acquisition, project administration, supervision and writing – original draft and writing – review & editing.

---

**Figure S1.**
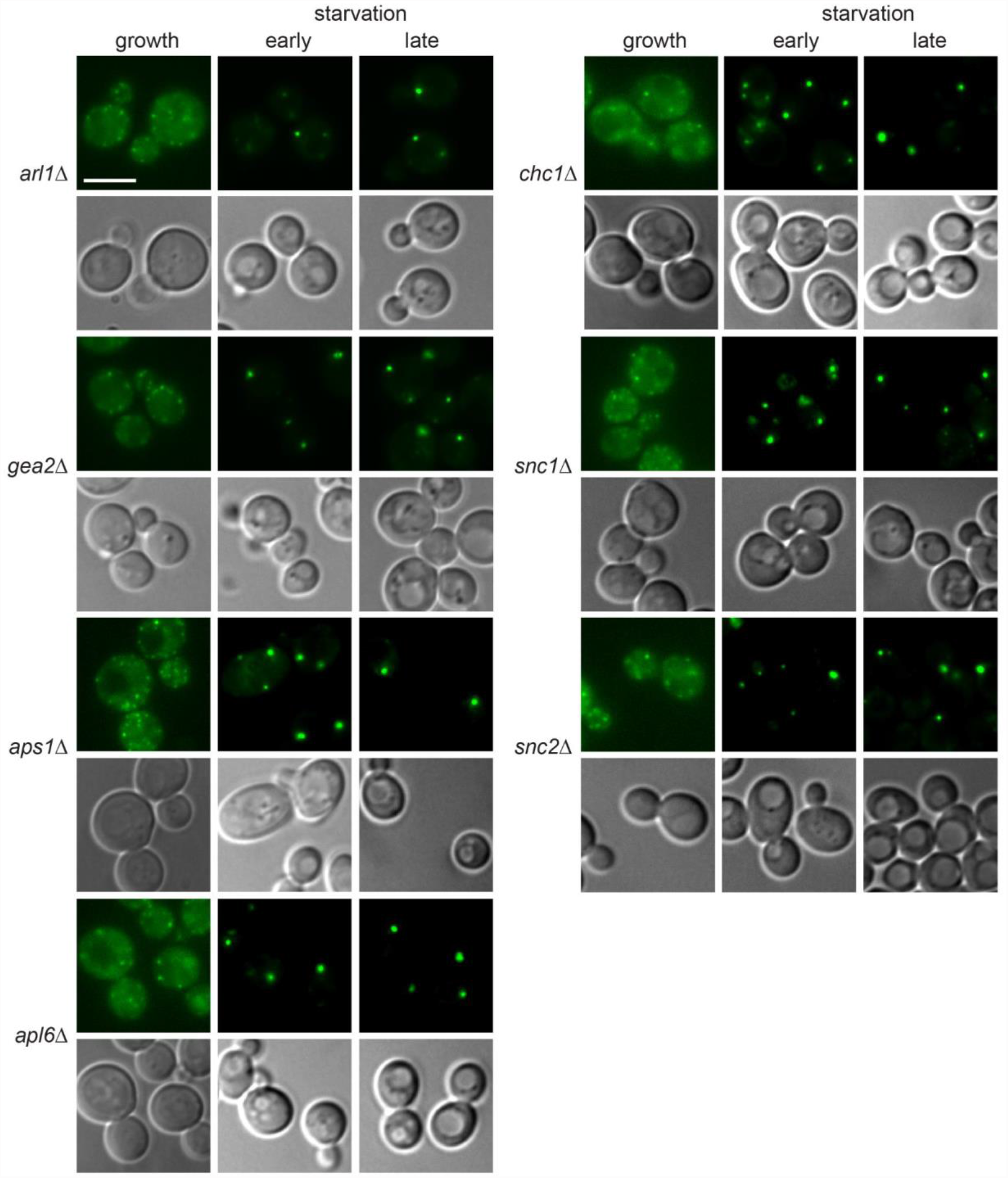
No CUPS defect in cells lacking Gea2, Arl1, Chc1, Apl6, Aps1, Snc1 or Snc2. The indicated deletion strains expressing Grh1-2xGFP were grown to log phase and starved for 2.5h. Scale bar = 2μm

**Figure S2.**
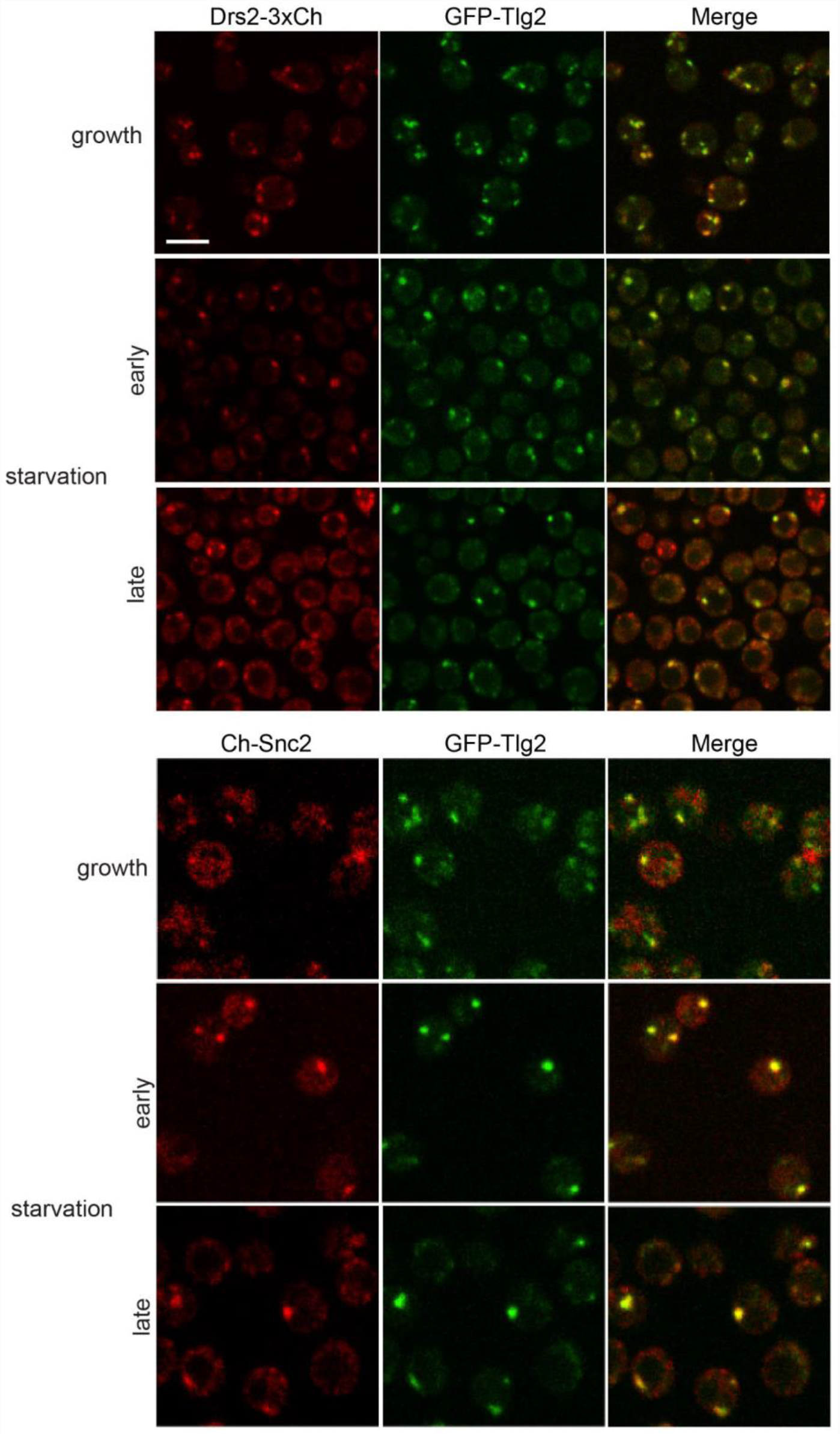
Drs2, Tlg2 and Snc2 label the same compartment in starvation – TCUPS. Cells co-expressing GFP-Tlg2 with Drs2-3xmCherry or mCherry-Snc2 were visualized by spinning disk confocal microscopy in the indicated conditions. In both combinations the average Pearson’s coefficient increased from ∼0.3 in growth to ∼0.7 in starvation (n=25-40 cells). Scale bar = 2μm.

**Figure S3.**
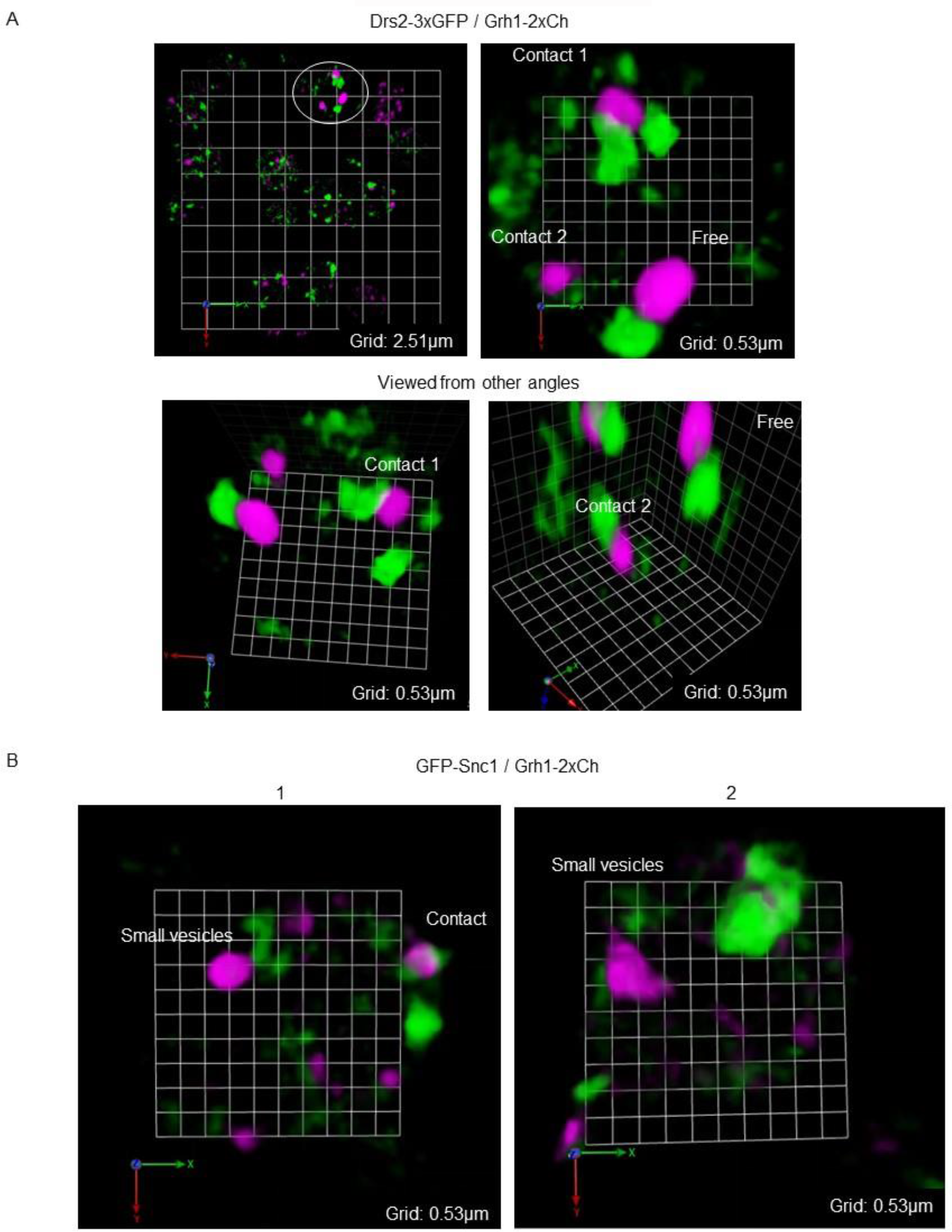
Drs2 and Snc2 also label small vesicles that contact with, or are near CUPS. Grh1-2xCh (magenta) cells co-expressing either (A) Drs2-3xGFP (green) or (B) GFP-Snc2 (green) were cultured in starvation condition for 1 hour. Movies S4-8; 10 sec intervals, 20x speed.

**Supplemental movies 1-3.** CUPS form by dynamic fusion and fission of existing membranes. Grh1-2xCherry (magenta) with GFP-Snc2 (green) at 1 hour of starvation. Movie S1 - smaller Grh1 structures can fuse. Movies S2 and S3 – larger Grh1 structures can separate and disperse. 10 sec intervals, 20x speed.

